# Identification and genetic diversity analysis of a male-sterile gene (*MS1*) in Japanese cedar (*Cryptomeria japonica* D. Don)

**DOI:** 10.1101/2020.05.09.085464

**Authors:** Yoichi Hasegawa, Saneyoshi Ueno, Fu-Jin Wei, Asako Matsumoto, Kentaro Uchiyama, Tokuko Ujino-Ihara, Tetsuji Hakamata, Takeshi Fujino, Masahiro Kasahara, Takahiro Bino, Katsushi Yamaguchi, Shuji Shigenobu, Yoshihiko Tsumura, Yoshinari Moriguchi

## Abstract

Identifying causative genes for a target trait in conifer reproduction is challenging for species lacking whole-genome sequences. In this study, we searched for the male-sterility gene (*MS1*) in *Cryptomeria japonica*, aiming to promote marker-assisted selection (MAS) of male-sterile *C. japonica* to reduce the pollinosis caused by pollen dispersal from artificial *C. japonica* forests in Japan. We searched for mRNA sequences expressed in male strobili and found the gene CJt020762, coding for a lipid transfer protein containing a 4-bp deletion specific to male-sterile individuals. We also found a 30-bp deletion by sequencing the entire gene of another individual with the *ms1*. All nine breeding materials with the allele *ms1* had either a 4-bp or 30-bp deletion in gene CJt020762, both of which are expected to result in faulty gene transcription and function. Furthermore, the 30-bp deletion was detected from three of five individuals in the Ishinomaki natural forest. From our findings, CJt020762 was considered to be the causative gene of *MS1*. Thus, by performing MAS using two deletion mutations as a DNA marker, it will be possible to find novel breeding materials of *C. japonica* with the allele *ms1* adapted to the unique environment of each region of the Japanese archipelago.

## Introduction

The breeding of forest trees is not as advanced as that of food crops and the planting trees are genetically close to individual trees of natural forests. Recent progress in genome analysis technology has promoted genome-based breeding techniques in tree plantations^1^. However, the construction of whole-genome sequences in conifers, which are important plantation trees, has been delayed due to the large size of their genome (>10 Gb) and extensive repetitive elements^2^. It is, therefore, challenging to identify the causative genes for a given target breeding trait in conifer species.

*Cryptomeria japonica* D. Don (Cupressaceae) covers over 4.5 million hectares, accounting for 44 % of all Japanese artificial forests^3^. As a result, in 2008, 26.5 % of Japanese residents had an allergy to *C. japonica* pollen^4^. Pollinosis caused by Japanese cedar, *C. japonica*, is a widespread social problem in Japan. Genetically male-sterile *C. japonica* trees are expected to play an important role in reducing the amount of dispersed pollen. In general, the frequency of male-sterile *C. japonica* trees is low within the Japanese population: for example, Igarashi *et al*. found two male-sterile trees among 8,700 trees in a 19 ha artificial forest^5^. To date, 23 genetically male-sterile *C. japonica* trees have been identified in Japan^6^. Based on the results of test crossings, four recessive male-sterile genes, *MS1*, *MS2*, *MS3 and MS4*, have been identified^6–9^ and they were mapped on linkage map^10^. Out of these genes, *MS1* is the most frequent in *C. japonica* male-sterility breeding materials, with 11 male-sterile trees homozygous for *MS1* (*ms1/ms1*) and five male-fertile trees heterozygous for *MS1* (*Ms1/ms1*) found in Japan^5,6,11–18^. The *C. japonica* trees that are homozygous for *ms1* become male-sterile due to the failure of exine development during microspore formation^16,19^, but it remains unclear whether *MS1* is caused by a single genetic mutation event. Additionally, there is no information about the geographical distribution of the genetic mutation responsible for *MS1*.

In recent years, the tightly linked DNA markers to perform marker-assisted selection (MAS) of individuals with *MS1* have been developed^20–25^. Mishima *et al*. suggested reCj19250 (the DEAD-box RNA helicase gene) as a candidate gene for *MS1* in *C. japonica*^23^. However, Ooi-7 heterozygous for *MS1* could not be selected with two SNP markers contained in reCj19250^22^, suggesting that reCj19250 was not the causative gene of *MS1*. Thus, to efficiently select individuals with *ms1* using MAS from various *C. japonica* populations, it is essential to search for genetic mutations that can consistently explain the *MS1* phenotype.

For the MAS of *C. japonica* trees that are homozygous or heterozygous for *MS1* and the breeding of male-sterile *C. japonica* trees adapted to the unique environments in each region of the Japanese archipelago, it is important to elucidate the diversity of genetic mutations responsible for *MS1* and the phylogenetic origin of *MS1*. Thus, in this study, we conducted the following analyses: (1) to identify the genetic mutation specific to homozygous or heterozygous individuals for *MS1* from mRNA sequences expressed in male strobili of *C. japonica*; (2) the functional annotation of candidate genes and deletion mutations causing *ms1*; and (3) construct the haplotype network and haplotype map of *MS1* candidate genes using breeding materials with *ms1* and trees from natural forests on the Japanese archipelago.

## Results

### RNA sequencing in the coding region of CJt020762

RNA-sequencing (RNA-Seq) data were analysed to identify the genetic variation specific to individuals with the recessive allele (*ms1*) of *MS1* among the cDNA sequences expressed in male strobili of *C. japonica*. RNA-Seq data from six libraries (Table S1) were used^26^. The genotype of *MS1* for these samples was determined based on phenotype and/or artificial crossing. cDNA sequences were screened using the following two criteria for *MS1* candidate genes: (1) sequences expressed in male strobili but not in needles and inner bark and (2) sequences with two-fold higher expression in male-fertile strobili compared to male-sterile strobili were selected.

From the above procedure, 88 cDNA sequences were obtained as candidate genes of *MS1*. One of them, CJt020762, contained an SNP marker (AX-174139329; Hasegawa *et al.*^22^) located in the same position (0cM) as *MS1* in the linkage map of the Fukushima-1 × Ooi-7 family. Furthermore, a 4-bp deletion causing a frameshift mutation was found in the coding region of CJt020762 (Figures 1 and 2). This deletion was not detected in wild-type homozygous individuals but was detected heterozygously in three heterozygous individuals for *MS1*, and homozygously in two male-sterile individuals. Therefore, the presence of the *ms1* allele and this genetic mutation in the CJt020762 region were consistent with *MS1*. We hereafter focused our analysis on this gene.

**Figure 1.**
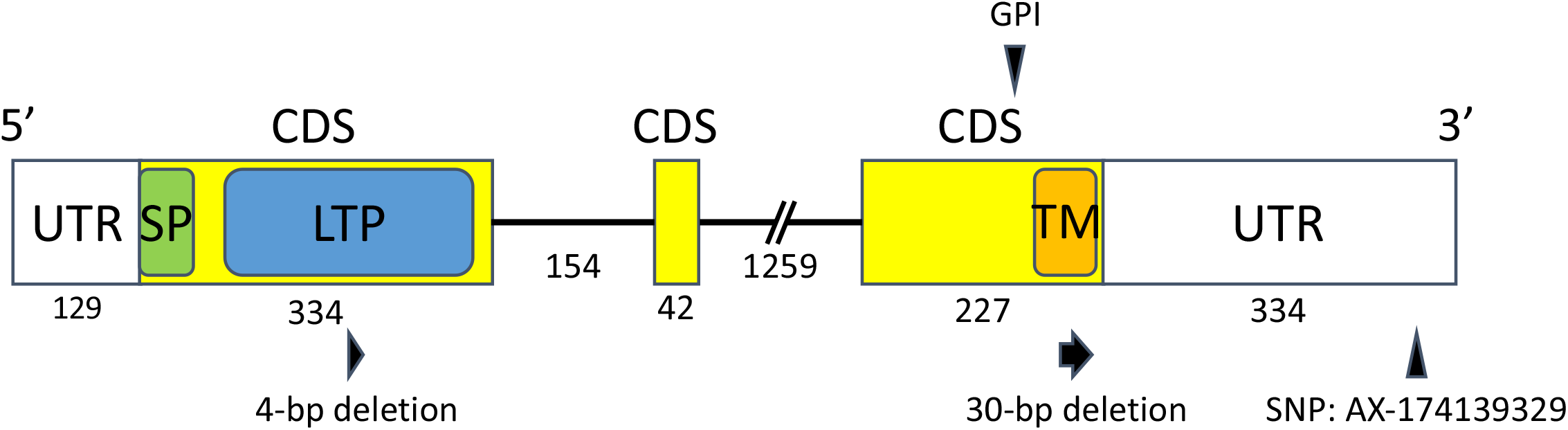
Schematic representation of CJt020762 including the two untranslated regions (UTRs), the three coding sequence (CDS) regions and two introns. The numbers indicate the length of nucleotide sequences in each region. LTP, predicted plant lipid transfer protein domain; SP, signal peptide; TM, transmembrane domain; GPI, potential modification site of GPI (glycosylphosphatidylinositol) anchor domain. Two deletion mutations were found on CJt020762. AX-174139329 indicates the SNP marker located at 0.0 cM from *MS1* on the linkage map constructed for the Fukushima-1×Ooi-7 family (Hasegawa *et al.*^22^).

**Figure 2.**
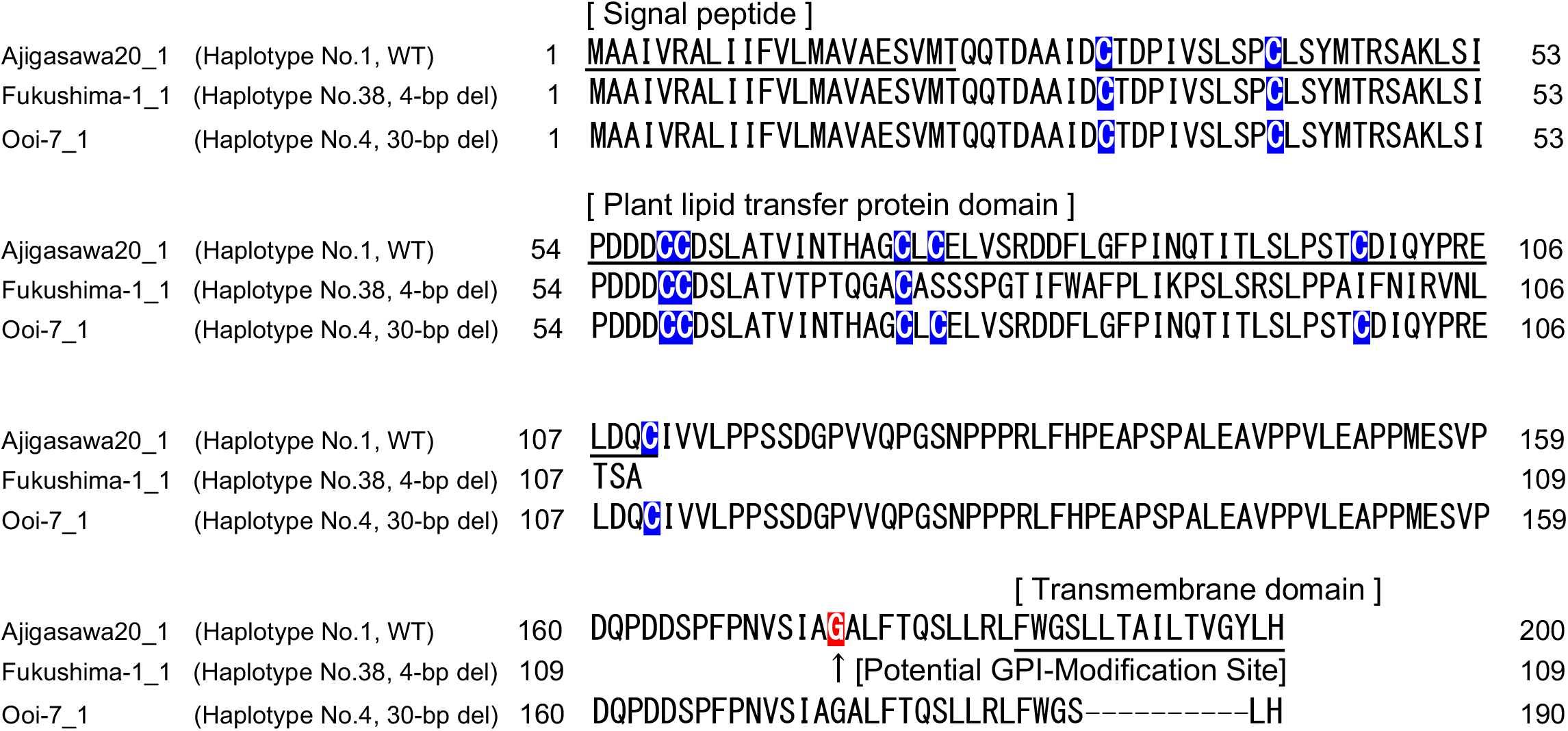
Amino-acid sequences for CJt020762. Underlines indicate the putative signal peptide, plant lipid transfer protein (LTP) domain and transmembrane domain. The blue boxes indicate the positions of eight conserved cysteine residues characteristic of LTPs. The red box indicates a potential modification site in the GPI (glycosylphosphatidylinositol) anchor domain.

CJt020762 codes for a lipid transfer protein gene containing a signal peptide, a plant lipid transfer protein domain, and a transmembrane domain in the coding region (Figure 1). Furthermore, the glycosylphosphatidylinositol (GPI) anchor domain that contributes to fixing the protein on the outside of the plasma membrane was predicted at the C-terminus of CJt020762 (Figures 1 and S1).

### Genomic DNA sequence of *MS1*

Eight primers were designed based on the cDNA sequence of CJt020762 (Table S2), and the Sanger method was used to sequence the genomic DNA of Fukushima-1 (Table 1). The genome sequence of CJt020762 was 2,556-bp long, containing three coding sequence (CDS) regions (Figures 1 and S2).

**Table 1.**
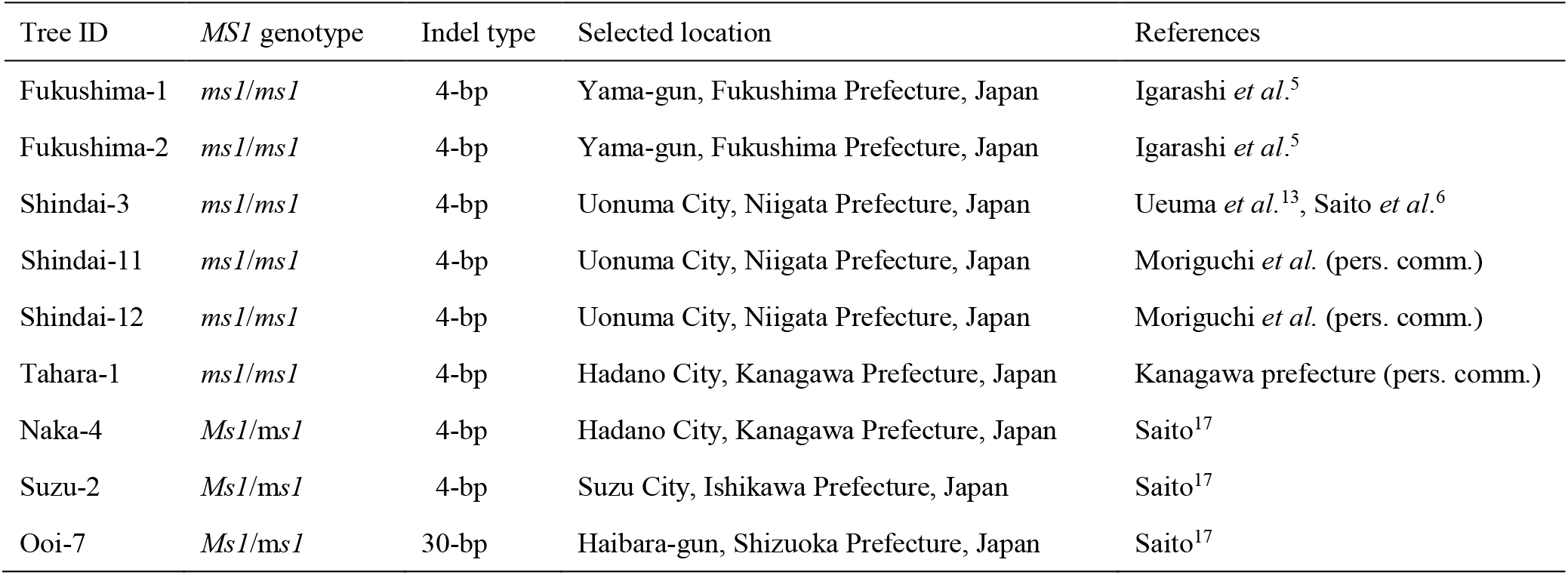
The breeding materials of *Cryptomeria japonica* with male-sterile gene 1 (*MS1*) used in this study.

### Haplotype composition of CJt020762 in breeding materials and individuals from natural forests

The haplotype sequences of CJt020762 of nine breeding materials with *ms1* and 74 individuals from 18 natural forest populations were determined. As a result, 49 haplotypes were detected from 83 individuals (Figure 3 and Table S3). Of these haplotypes, only two (No. 38 (*ms1-1*) and No. 4 (*ms1-2*)) contained the deletion in the coding region. In haplotype No. 38, the frameshift mutation caused by the 4-bp deletion occurred in the middle of the plant lipid transfer protein domain (Figure 2). On the other hand, in haplotype No. 4, the deletion of amino-acids occurred in the transmembrane domain due to the 30-bp deletion (Figure 2). Furthermore, the modification site of the GPI anchor domain on the C-terminal side, which is important for transmembrane functioning, was lost after the 30-bp deletion (Figure S1). All individuals homozygous for *MS1* (*ms1*/*ms1*) (i.e., Fukushima-1, Fukushima-2, Shindai-3, Shindai-11, Shindai-12, and Tahara-1) had the homozygous haplotype No. 38 (Figure 3 and Table 1), while individuals heterozygous for *MS1* (*Ms1*/*ms1*) shown in Table 1 were heterozygous with haplotype No. 38 (Naka-4, Suzu-2) or No. 4 (Ooi-7) (Figure 3 and Table 1). Haplotype No. 38 was absent from the 74 individuals from natural populations (Figure 3 and Table S3). On the other hand, 17 individuals with haplotype No. 1, which is considered to be the ancestor of haplotype No. 38 (*ms1-1*), were found in 13 natural populations (Ajigasawa, Mizusawa, Nibetsu, Yamanouchi, Donden, Bijodaira, Shimowada, Ashu, Tsuyama, Wakasugi, Azouji, Oninome and Yakushima; Figures 3 and 4, Table S3). Haplotype No. 4 (*ms1-2*) with the 30-bp deletion was found in three individuals from the Ishinomaki natural population (Figures 3 and 4, Table S3). Furthermore, five individuals with haplotype No. 2, which is considered to be the ancestor of haplotype No. 4, were found in three natural populations (Oki, Azouji and Oninome; Figures 3 and 4, Table S3).

**Figure 3.**
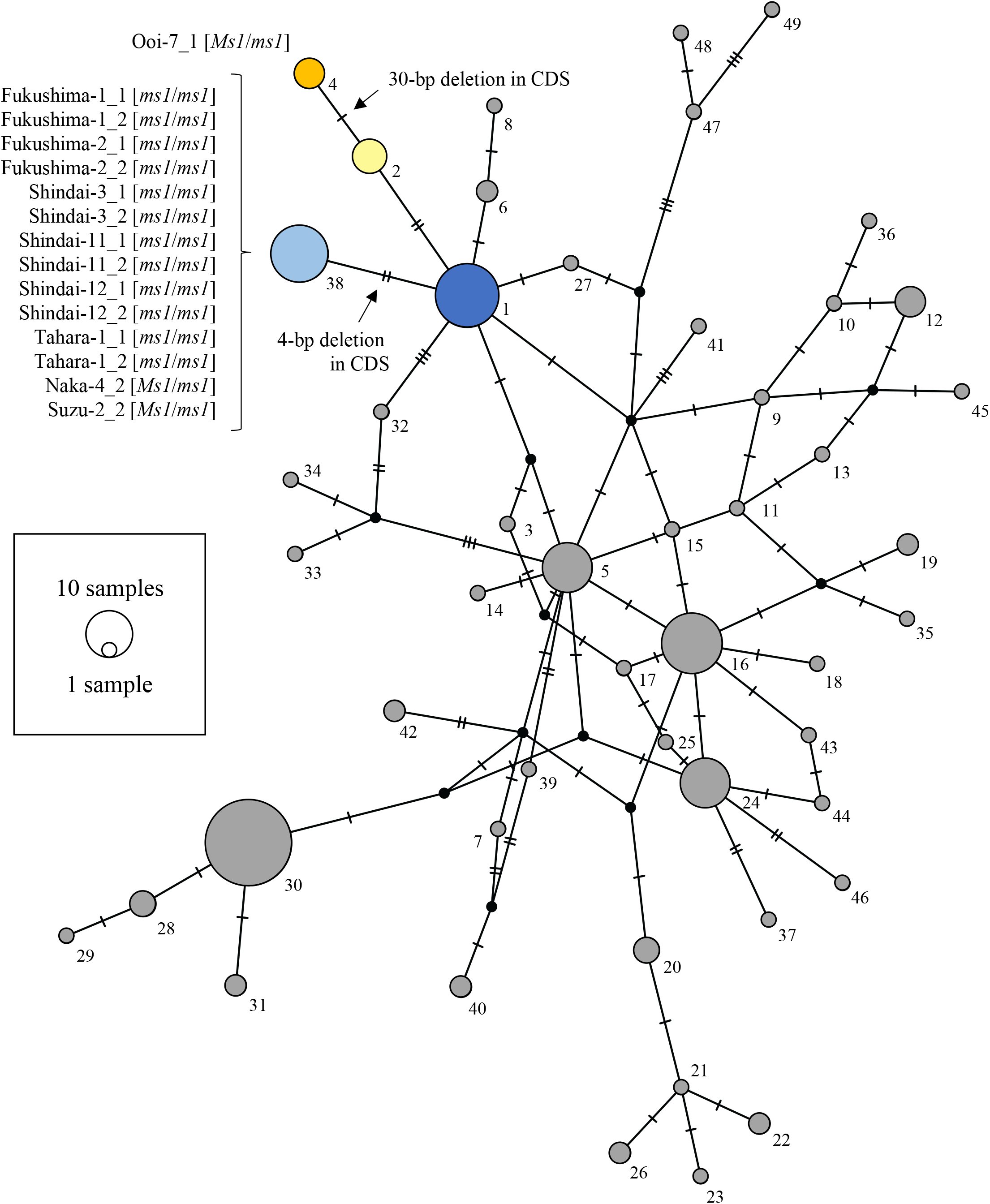
The haplotype network of CJt020762, based on 49 haplotypes determined from the 83 individuals including nine breeding materials with *ms1*. Numbers indicate the haplotype numbers corresponding to Table S3. Haplotype names (e.g. Fukushima-1_1) indicate the haplotype of breeding materials with *ms1*. [*ms1/ms1*] indicates homozygozity for *MS1*, and [*Ms1/ms1*] indicates heterozygozity for *MS1*. The node represents a haplotype. Node size represents haplotype frequency. The light blue segment represents the haplotype with a 4-bp deletion (No. 38); the blue segment represents the ancestor haplotype of the haplotype No. 38 (No. 1); the orange segment represents the haplotype with the 30-bp deletion (No. 4); the yellow segment represents the ancestor haplotype of the haplotype No. 4 (No. 2); and the grey segment represents the remaining 45 haplotypes. The horizontal line between nodes represents one SNP or one insertion/deletion (indel).

**Figure 4.**
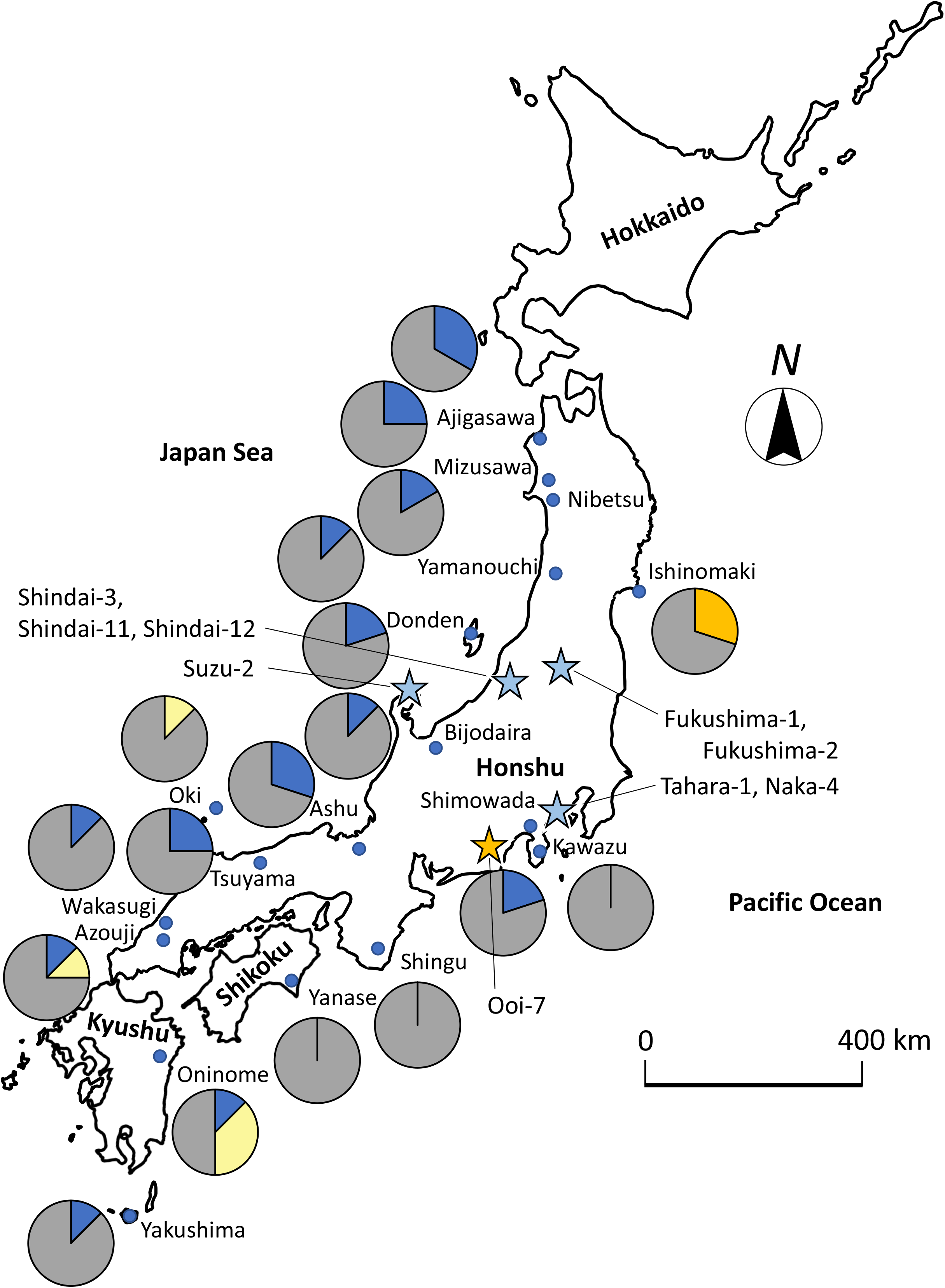
Geographical distribution of genomic DNA haplotypes of CJt020762 and breeding materials with *ms1*. The light blue stars pinpoint locations where *ms1* breeding materials with a 4-bp deletion were selected; the orange star represents the location where Ooi-7 with a 30-bp deletion was selected; the blue segment represents the ancestor haplotype of the haplotype with a 4-bp deletion (No. 1); the orange segment represents the haplotype with a 30-bp deletion (No. 4); the yellow segment represents the ancestor haplotype of the haplotype with the 30-bp deletion (No. 2); and the grey segment represents the remaining 45 haplotypes. All haplotype No. correspond to those in Figure 3 and Table S3.

Amino-acid substitutions were found in six haplotypes from natural populations (Haplotype No. 36, 37, 39, 40, 41 and 42; Table S4). However, they were not detected in the breeding materials for *ms1*.

## Discussion

In this study, we searched for genetic mutations specific to *C. japonica* breeding materials with *ms1* using RNA-Seq data. Based on the results, we identified CJt020762 as a causative gene for *MS1*. There is additional existing evidence supporting this finding. Firstly, CJt020762 contains the SNP marker (AX-174139329) mapped on 0 cM from *MS1* in the linkage map of the Fukushima-1 × Ooi-7 family^22^, indicating that CJt020762 was a gene in the vicinity of *MS1*. Secondly, CJt020762 was expressed in male strobili, where it is active. Thirdly, both 4-bp and 30-bp deletions are expected to cause dysfunction in this particular gene: the 4-bp deletion mutation deleted the amino-acid sequence of the plant lipid transfer protein domain, causing a frameshift, while the 30-bp deletion mutation deleted 10 (63 %) amino-acids in the transmembrane domain (16 amino-acids). Furthermore, the modification site of the GPI anchor domain that contributes to fixing the protein to the outside of the plasma membrane was lost after the 30-bp deletion. These amino-acid sequence mutations are likely to result in the malfunction of the lipid transfer protein coded by CJt020762. The fourth piece of evidence suggesting that CJt020762 is the causative gene of *MS1* comes from the mapping family Fukushima-1 × Ooi-7, where male-sterility occurred in one-half of offsprings^22^. Result of the DNA sequencing (Table S3) revealed that Fukushima-1 was homozygous for the 4-bp deletion haplotype (*ms1-1*/*ms1-1*) and that Ooi-7 was heterozygous, with the 30-bp deletion and wild-type haplotype (*Ms1*/*ms1-2*). Therefore, it was clarified that even if CJt020762 is heterozygous for the 4-bp deletion haplotype and 30-bp deletion haplotype (*ms1-1*/*ms1-2*), male-sterility occurs. Such a phenomenon occurs when CJt020762 codes a protein essential for pollen production and the protein function is lost due to the 4-bp deletion in the plant lipid transfer protein domain and the 30-bp deletion in the transmembrane domain. This result also indicates that both the plant lipid transfer protein domain and the transmembrane domain contained in CJt020762 is essential for pollen production in *C. japonica*. The fifth and final supporting evidence stems from the functional and structural similarity of CJt020762 with wheat male-sterility genes. In wheat male-sterility (*Triticum aestivum*), both *ms1* and *ms5* possess recessive inheritance and single locus control, hence male-sterility is caused by the failure of exine development during microspore formation^27–30^. *MS1* in *C. japonica* and *ms1* and *ms5* in wheat have similar male-sterility phenotypes. In terms of protein structure, CJt020762 contains the signal peptide, plant lipid transfer protein domain, transmembrane domain, and the GPI anchor domain modification sites (Figure 1). This structure is similar to *ms1* and *ms5* in wheat^28–30^. Furthermore, while the plant lipid transfer protein domain and transmembrane domain of CJt020762 may be necessary for pollen production in *C. japonica*, an analysis of mutant wheat revealed that both the lipid transfer protein domain and transmembrane domain of wheat *ms1* were necessary for pollen production^29^. Because of the similarities with wheat male-sterility genes, CJt020762 is thought to be essential for pollen production, as well as *ms1* and *ms5* in wheat, despite their respective amino-acid sequences being different to each other, with percentage identity ranging from 27.0 to 31.4 for the highest scoring segment pairs in BLASTP.

Based on above evidence, we propose that the CJt020762 section is the *MS1* gene itself. We clarified the genomic DNA haplotypes of CJt020762 in 83 individuals, including nine breeding materials with *ms1* and 74 individuals from 18 natural forests. Eight individuals (Fukushima-1, Fukushima-2, Shindai-3, Shindai-11, Shindai-12, Tahara-1, Naka-4, Suzu-2) with *ms1-1* (haplotype No. 38) had a 4-bp deletion in the plant lipid transfer protein domain. Additionally, Ooi-7 with *ms1-2* (haplotype No. 4) had a 30-bp deletion in the transmembrane domain. These results suggest that these deletions were caused by independent two genetic mutation events. In the Ishinomaki natural forest, haplotype No. 4 with the 30-bp deletion (*ms1-2*) was found in 3/5 individuals. This result indicates that individuals with haplotype No. 4 may be more frequent in the Ishinomaki natural forest. The distance between the selected location of Shindai-3, Shindai-11, Shindai-12 and that of Tahara-1, Naka-4 was approximately 300 km and that between the selected location of Ooi-7 and the Ishinomaki natural forest was approximately 450 km. Therefore, the single genetic deletion mutations were dispersed over hundreds of kilometers. Furthermore, haplotype No. 4 (*ms1*-*2*) was derived from ancestral haplotype No. 2, which is distributed in three southern populations (Oki, Azouji and Oninome). As haplotype No. 4 was found on the Pacific Ocean side in the central and northern regions of Honshu island (Ooi-7 and Ishinomaki forest), it may have expanded further north along the Pacific side. Using markers developed during this study, a number of individuals with *ms1* can be selected from all over Japan in the future. Identifying the CJt020762 haplotype of these individuals and comparing it to the distribution of ancestral haplotypes will clarify the historical gene flow of *ms1*.

The findings of this study suggest that CJt020762 is the causative gene for *MS1* of *C. japonica* and contribute to the MAS of *MS1* for breeding of male-sterile trees. Although the 16 individual trees with *ms1* have – until now – all been found in Japan^5,11–14,16–18^, it is necessary to select more individuals by MAS to obtain breeding materials in each region, a *C. japonica* had differentially adapted to unique environments in different regions of the Japanese archipelago, such as around the Japan Sea (heavy snow in winter) or along the Pacific coast (dry in winter)^31^. To definitely prove that CJt020762 is the causative gene for *MS1*, it is essential to show that male-sterility occurs by knocking out CJt020762 in *C. japonica* through genome editing in future research.

## Methods

### RNA-sequencing and *MS1* annotation

To identify causative single nucleotide variations (SNVs) and/or insertions or deletions (indels) in expressed genes, we analysed RNA-sequencing (RNA-Seq) data from six libraries (Table S1) from the mapping family of *MS1*. These data have been reported in Ueno *et al.*^25^ and Wei *et al.*^26^. In this study, we focused on the levels of gene expression and selected candidate genes using the following criteria: 1) expression levels (in transcripts per million [tpm]) in leaf or bark tissue < 10; 2) expression levels (tpm) in fertile strobili was more than twice compared to those in sterile strobili. We selected 88 candidate genes for *MS1*, which were then examined for SNVs and indels. All of the read mapping of the RNA-Seq was checked manually using an IGV (Integrative Genome Viewer)^32^. After we identified the causative mutation for *MS1* from the RNA-Seq data, we examined the translated amino-acid sequences of the candidate gene (CJt020762) for the functional domain and annotated using SignalP-5.0^33^, SMART (https://smart.embl.de/), PRED-TMBB2^34^, and big-PI Plant Predictor^35^.

### Plant materials and DNA extraction

This study used six breeding materials (trees) homozygous for *MS1* (*ms1/ms1*), three breeding materials heterozygous for *MS1* (*MS1*/*ms1*), and 80 trees from 18 natural forests (Tables 1 and S5). Needle tissue was collected from all 89 trees; in natural forests, it was collected from an established scion garden through cutting. Genomic DNA was extracted from these needles using a modified CTAB method^36^.

### Sanger sequencing of CJt020762

PCR was performed in 15 μL volume containing approximately 60 ng of genomic DNA of Fukushima-1 (Table 1), 1× Multiplex PCR Master Mix (Qiagen, Hilden, Germany), and 0.2 μM of Primer_F3 and Primer_R (Figure S2 and Table S2). The thermal profile for the PCR was as follows: an initial denaturing step of 15 min at 95 °C, followed by 40 cycles of 30 s at 94 °C, 90 s at 63 °C and 90 s at 72 °C, before a final elongation step at 72 °C for 10 min using a GeneAmp 9700 PCR System (Applied Biosystems, PE Corp., Foster City, CA, USA). PCR products were purified using ExoSAP-IT (Affymetrix, Inc., Santa Clara, CA, USA). Direct sequencing was performed using the ABI PRISM BigDye Terminator version 3.1 Cycle Sequencing Kit (Applied Biosystems) on an ABI 3130 Genetic Analyser (Applied Biosystems). For sequencing, we also used additional internal primers (Primer_F2, Primer_F2-2, Primer_R2, Primer_R2-2, Primer_R2-3, Primer_R2-4) developed in this study (Figure S2 and Table S2). Sequence alignment was performed using CodonCode Aligner v.8.0.2 software (CodonCode Corporation, Dedham, MA, USA), followed by manual editing.

### Amplicon sequencing of CJt020762

We developed seven primer pairs using the PCR suite software^37^ (Figure S2 and Table S2) using the genomic sequence of CJt020762 for Fukushima-1, following the Sanger method. We used these primer pairs and 89 individual trees (Tables 1 and S5) for the sequencing of whole CJt020762. Single-plex PCR was performed in 10μL volume containing approximately 30 ng of genomic DNA, 1× Multiplex PCR Master Mix (Qiagen) and 0.2 μM of each primer pair. The thermal profile for PCR was as follows: an initial denaturing step of 15 min at 95 °C, followed by 35 cycles of 30 s at 94 °C, 90 s at 66 °C and 90 s at 72 °C before a final elongation step at 72 °C for 10 min by using a GeneAmp 9700 PCR System (Applied Biosystems). PCR products were pooled in equal volumes for each individual tree. Tag sequences for identifying individual trees were attached to the DNA fragments using the Access Array Barcode Library for Illumina Sequencers-384, Single Direction (Fluidigm Corporation, South San Francisco, CA, USA) and KAPA HiFi HotStart ReadyMix PCR Kit (KAPA BioSystems, Wilmington, MA, USA). PCR was performed in a 20 μL volume containing 2 μL of 20-times diluted PCR products, 1× KAPA HiFi HS ReadyMix and 4 μL of Access Array primers. The thermal profile for PCR was as follows: an initial denaturing step for 5 min at 95 °C, followed by 12 cycles of 15 s at 95 °C, 30 s at 60°C and 60 s at 72°C before a final elongation step at 72 °C for 3 min by using a GeneAmp 9700 PCR System (Applied Biosystems). After PCR, each DNA sample was purified using AMPure XP (Beckman Coulter, Brea, CA, USA). DNA concentration was determined using the Qubit fluorometer in the Qubit dsDNA HS Assay Kit (Thermo Fisher Scientific, Waltham, MA, USA). An equal amount of DNA from each sample was mixed to construct an amplicon sequencing library. The library was size-selected using BluePippin (1.5 % agarose cartridge, Sage Science, Beverly, MA, USA) under the range of 400-700 bp. The DNA concentration of the library was determined using the LightCycler 480 Real-Time PCR System (Roche, Basel, Switzerland) with KAPA Library Quantification Kit (KAPA Biosystems). Finally, the library was sequenced in 2×251-bp paired ends on MiSeq (Illumina, San Diego, CA, USA). Reads from each individual were automatically classified on MiSeq according to the tag sequences (DRR206539-DRR206627). After reads were cleaned using the Trimmomatic programme^38^, they were mapped onto the genomic sequence of CJt020762 from Fukushima-1 using the BWA mem algorithm^39^. Each mapping file (BAM) was imported into the Geneious software (Biomatters Ltd., Auckland, New Zealand) and possible SNVs sites were manually checked. The consensus sequence was exported, then haplotype sequences of CJt020762 were estimated using the Phase software^40,41^. Furthermore, to clarify phylogenetic relationships among the CJt020762 haplotypes, haplotype networks were created using the PopART software^42^. Six individual trees (Ajigasawa02, Bijodaira03, Tsuyama03, Tsuyama07, Yakushima02, and Yamanouchi04) were excluded from the haplotype analysis as insufficient DNA sequences were obtained.

## Acknowledgements

The authors thank Yasuyuki Komatsu and Nozomi Ohmiya for their assistance with laboratory work, Yukiko Ito, Hiroshi Saito, and Akira Ogura for providing breeding materials for this study. This work was supported by research grants #201421 from the Forestry and Forest Products Research Institute, Grant-in-Aid from the Program for Promotion of Basic and Applied Researches for Innovations in Bio-oriented Industry (No.28013B), the project of the NARO Bio-oriented Technology Research Advancement Institution (Research program on development of innovative technology, No.28013BC), and NIBB Collaborative Research Program (16-403, 17-405 and 18-408). Computations were partially performed on the supercomputer of AFFRIT, MAFF, Japan.

## Author Contributions

Y.H., S.U., A.M., K.U., T.U., M.K., S.S., Y.T. and Y.M. conceived and designed the experiments. Y.H., S.U., A.M., T.H., T.B. and K.Y. and Y.M. performed the experiments. Y.H., S.U., F.W., T.F., M.K., S.S. and Y.M. analysed the data. Y.H., S.U. and Y.M. wrote the paper.

## Additional Information

Figure S1. Results of a 30-bp deletion on the loss of potential GPI-modification sites. Peptide sequences of Ajigasawa20_1 and Ooi-7_1 were tested for prediction of potential GPI-modification site using big-PI Plant Predictor (Eisenhaber *et al.* 2003).

Figure S2. Genomic DNA sequence of CJt020762 for Fukushima-1 and primer site position.

Table S1. RNA-Seq libraries of *C. japonica* used to identify specific mutations in individuals with the male-sterility gene (*MS1*).

Table S2. Primer sequences used in this study. The primer sequences for amplicon sequencing contained either the CS1 adapter: ACACTGACGACATGGTTCTACA or CS2 adapter: TACGGTAGCAGAGACTTGGTCT for Access Array primers of Access Array Barcode Library for Illumina Sequencers-384, Single Direction (Fluidigm Corporation, South San Francisco, CA, USA).

Table S3. List of mutations in the CJt020762 genomic sequence.

Table S4. Amino-acid mutations in CJt020762. The amino-acid sequence of Ajigasawa20_1 was used as a reference haplotype (Haplotype No. 1).

Table S5. Summary of the *C. japonica* trees collected from natural forests for use in amplicon sequencing of CJt020762.

