## Supplementary material for "Identification and genetic diversity analysis of a male-sterile gene (*MS1*) in Japanese cedar (*Cryptomeria japonica* D. Don)": Figure S1

| Haplotypes | Wild Type Ajigasawa20_1 | *ms1* Ooi-7_1 (30bp deletion ) |
|---|---|---|
| Peptide sequence | MAAIVRALIIFVLMAVAESVMTQQTDAAIDCTDPIVSLSPCLSYMTRSAKLSIPDDDC<br>CDSLATVINTHAGCLCELVSRDDFLGFPINQTITLSLPSTCDIQYPRELDQCIVVLPPSS<br>DGPVVQPGSNPPPRLFHPEAPSPALEAVPPVLEAPPMESVPDQPDDSPFPNVSIAG<br>ALFTQSLLRLFWGSLLTAILTVGYLH | MAAIVRALIIFVLMAVAESVMTQQTDAAIDCTDPIVSLSPCLSYMTRSAKLSIPDDDC<br>CDSLATVINTHAGCLCELVSRDDFLGFPINQTITLSLPSTCDIQYPRELDQCIVVLPPSS<br>DGPVVQPGSNPPPRLFHPEAPSPALEAVPPVLEAPPMESVPDQPDDSPFPNVSIAG<br>ALFTQSLLRLFWGSLH |
| Prediction of potential C-terminal GPi-modification site | Potential GPI-modification site was found.<br>Quality of the site ............... : S<br>Sequence position of the omega-site : **176**<br>Score of the best site ........... : -3.28 (PValue = 2.791270e-03) | **None** potential GPI-modification site was found.<br>Among all positions checked, sequence position 160 had the best score. |

Figure S1 Results of a 30-bp deletion on the loss of potential GPI-modification sites. Peptide sequences of Ajigasawa20_1 and Ooi-7_1 were tested for prediction of potential GPI-modification site using big-PI Plant Predictor (Eisenhaber *et al.* 2003).
