## Supplementary figures and images for "Identification and genetic diversity analysis of a male-sterile gene (*MS1*) in Japanese cedar (*Cryptomeria japonica* D. Don)"

### Figure S2

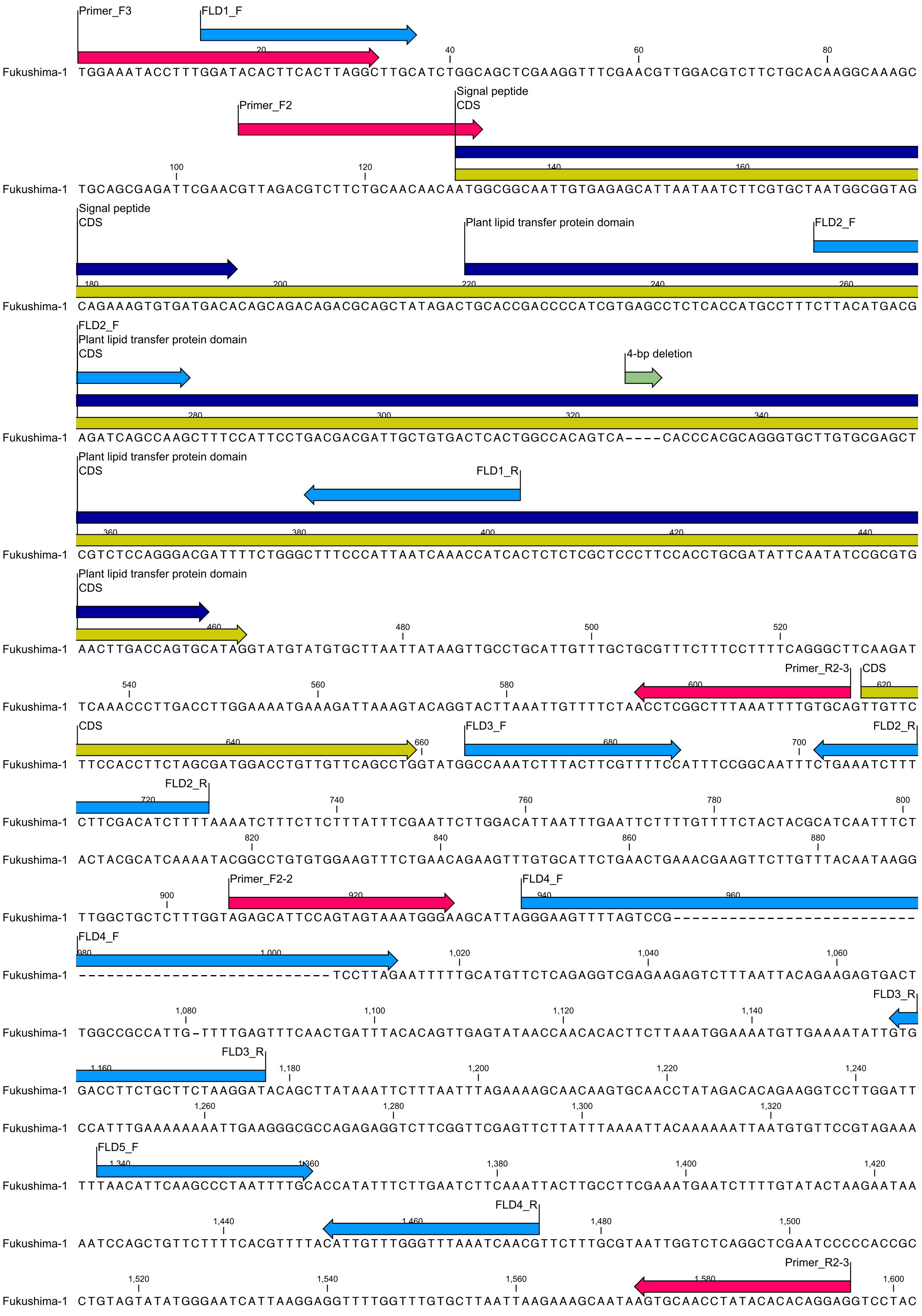




Figure S2 Genomic DNA sequence of CJt020762 for Fukushima-1 and primer site position.

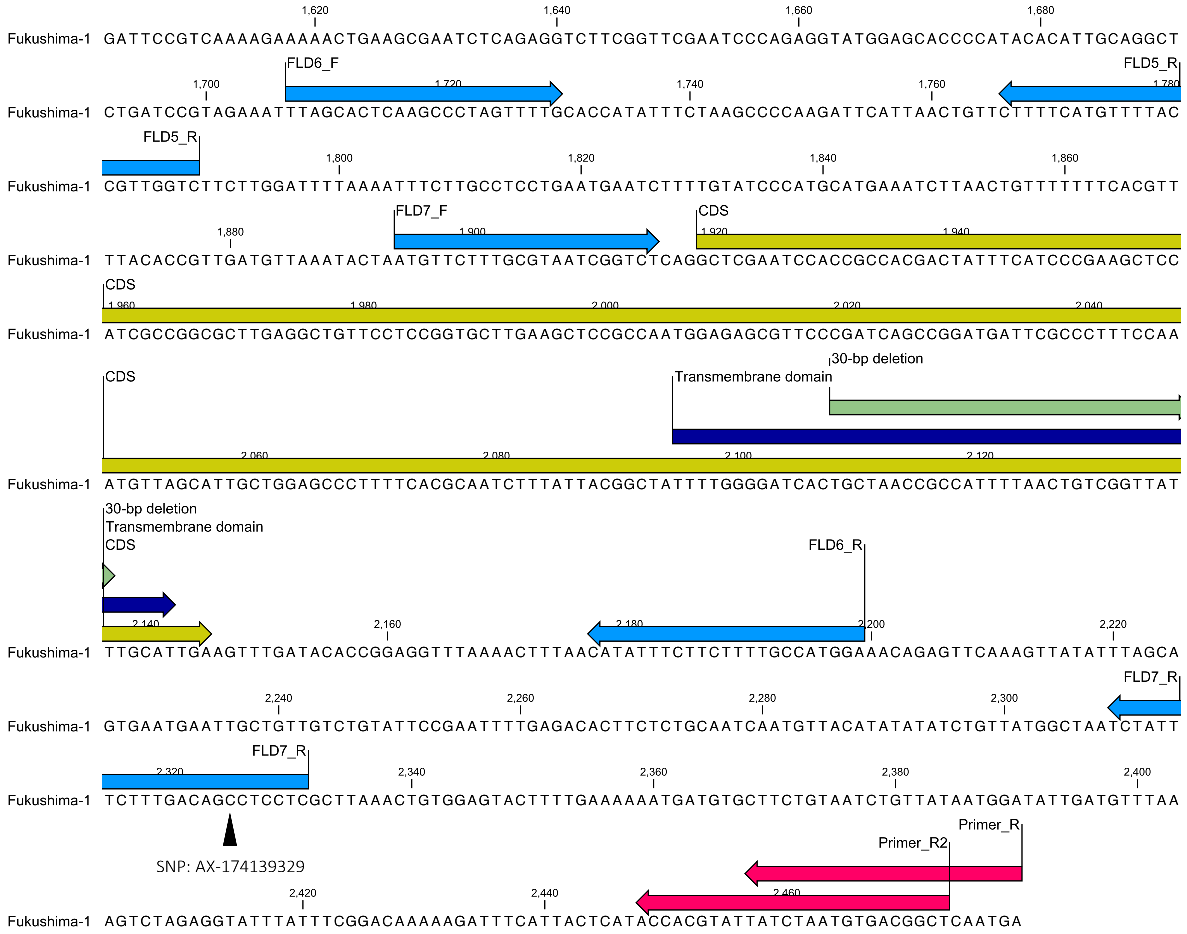


Figure S2 (continued)
